# Matrix porosity is associated with *Staphylococcus aureus* biofilm survival during prosthetic joint infection

**DOI:** 10.1101/2024.12.06.627279

**Authors:** Mohini Bhattacharya, Tyler D. Scherr, Jessica Lister, Tammy Kielian, Alexander R. Horswill

## Abstract

Biofilms are a cause of chronic, non-healing infections. *Staphylococcus aureus* is a proficient biofilm forming pathogen commonly isolated from prosthetic joint infections that develop following primary arthroplasty. Extracellular adhesion protein (Eap), previously characterized in planktonic or non-biofilm populations as being an adhesin and immune evasion factor, was recently identified in the exoproteome of *S. aureus* biofilms. This work demonstrates that Eap and its two functionally orphaned homologs EapH1 and EapH2, contribute to biofilm structure and prevent macrophage invasion and phagocytosis into these communities. Biofilms unable to express Eap proteins demonstrated increased porosity and reduced biomass. We describe a role for Eap proteins *in vivo* using a mouse model of *S. aureus* prosthetic joint infection. Results suggest that the protection conferred to biofilms by Eap proteins is a function of biofilm structural stability that interferes with the leukocyte response to biofilm-associated bacteria.

## Introduction

The Gram-positive pathogen, *Staphylococcus aureus,* has been the causative agent of >119,000 bloodstream infections in the United States, with nearly 20,000 deaths caused by <u>m</u>ethicillin <u>r</u>esistant *<u>S</u>. <u>a</u>ureus* (MRSA) [1], [2]. Recent studies show that while measures to control hospital-associated bacterial transmission have reduced the occurrence of serious *S. aureus* infections, this success has been slowing [2]. Approximately 1-3% of total hip and knee arthroplasties continue to be complicated by infection, resulting in longer hospital stays, higher occurrence of revision surgeries, and decreased 5-year survival rates [3], [4]. Along with the acquisition of resistance to many currently prescribed antibiotics, the ability of *S. aureus* to form biofilms during chronic infections has made this pathogen a substantial cause of concern with approximately 20% of surgical site infections reported to be associated with *S. aureus* [4], [5]. Biofilm-associated bacteria can tolerate up to 1,000 times the antibiotic concentrations that are found to be effective against planktonic or non-biofilm forms of the same strain [6], [7]. Furthermore, biofilms are generally recognized as a distinct lifestyle with uniquely attributable virulence mechanisms [8], [9], [10], [11]. Bacteria communicate within biofilms via quorum sensing molecules that allow for the development of shared, public goods [12]. A consequence of this is the formation of a protective matrix surrounding the biofilm, consisting of one or more components, including proteins, DNA and/or polysaccharides [13], [14]. Since biofilms are formed under nutritional or environmental stresses, this often allows the pathogen to evade host antimicrobial responses until conditions that are favorable for planktonic growth become available [15]. When this occurs, biofilm-associated bacteria disperse from the community and often cause disseminated infections including, but not limited to, serious bloodstream-associated conditions [15], [16], [17]. Therefore, it is imperative that biofilm-associated phenotypes are considered in the prevention of persistent *S. aureus* infections [18].

One of the major secreted and surface-associated proteins found in the *S. aureus* biofilm matrix is <u>e</u>xtracellular <u>a</u>dherence <u>p</u>rotein (Eap), which is reported to ubiquitously bind to numerous host proteins as well as bacterial and host DNA [19]. Eap is primarily secreted from *S. aureus* but is also described as being able to subsequently bind to the bacterial surface via the activity of a neutral phosphatase and other, yet uncharacterized factors [20]. Previous studies with planktonic bacteria attribute important anti-inflammatory and anti-angiogenic properties to Eap during *S. aureus* endovascular infection [21]. While this protein has been demonstrated to contribute to biofilm formation under conditions of stress, including iron starvation and the presence of serum, the role that Eap could play as a virulence factor during biofilm growth, is currently understudied [22], [23]. *S. aureus* also expresses two functional orphans of Eap, EapH1 and EapH2, recently reported to protect the bacterium against neutrophil-derived proteases [24], [25]. Macrophages have been established as being crucial for an effective immune response to *S. aureus* in wounds and foreign body-associated infections [26], [27], [28], [29]. Here we show that the expression of the three Eap proteins (Eap, EapH1 and EapH2) prevents macrophages from invading and phagocytosing *S. aureus* biofilm bacteria. These phenotypes are specific to macrophages since neutrophils were relatively unaffected by the presence of Eap. Additionally, using an established murine model of prosthetic joint infection we show that the inability to express Eap causes a significant reduction in bacterial burdens in the joint as well as surrounding tissue [26], [30]. Together these data provide evidence for the role of Eap as a biofilm structural protein that promotes *S. aureus* orthopedic infections.

## Results

### Eap proteins contribute to biofilm biomass and structure

To understand if Eap plays a role in biofilm development, we compared the gross biofilm biomass of the most commonly isolated *S. aureus* lineage, USA300 (hereafter referred to as WT) to an isogenic mutant lacking *eap* as well as its two functionally orphaned homologs, *eapH1* and *eapH2* (hereafter *Δeap*) using an established crystal violet-based assay [31]. Biomass comparisons of biofilms from both strains grown for 24 hours showed that *Δeap* bacteria have a significant loss of biomass compared to WT biofilms (**Figure 1A**). The immunomodulatory protein IsaB is another DNA binding protein that is abundantly expressed as part of the biofilm exoproteome of common clinical strains of *S. aureus* [32], [33]. We therefore generated a mutant of the *isaB* gene in the Δ*eap* strain background. Bacteria lacking IsaB in addition to the three Eap proteins, formed biofilms with biomass comparable to *Δeap* bacteria (**Figure 1A**). These results indicate that while the 3 Eap proteins are important for biofilm structure, the immunomodulatory surface protein IsaB does not significantly contribute to *in vitro* biofilm formation under these conditions. Confocal microscopy was used to further investigate the differences in gross biomass of 24-hour biofilms observed with crystal violet assays. 3D images indicated that when compared to WT biofilms, *Δeap* and *ΔeapΔisaB* biofilms showed a loss of structure and thickness (**Figure 1B**). Quantification of biofilm biomass from confocal microscopy confirmed that *Δeap* biofilms have a significant loss of thickness compared to WT biofilms and that there was no further decreases in the isogenic *ΔeapΔisaB* strain (**Figure 1C**). Collectively these data indicate that Eap proteins contribute to the gross biofilm biomass and that these proteins may also play a specific role in the overall structure of *S. aureus* biofilms.

**Figure 1.**
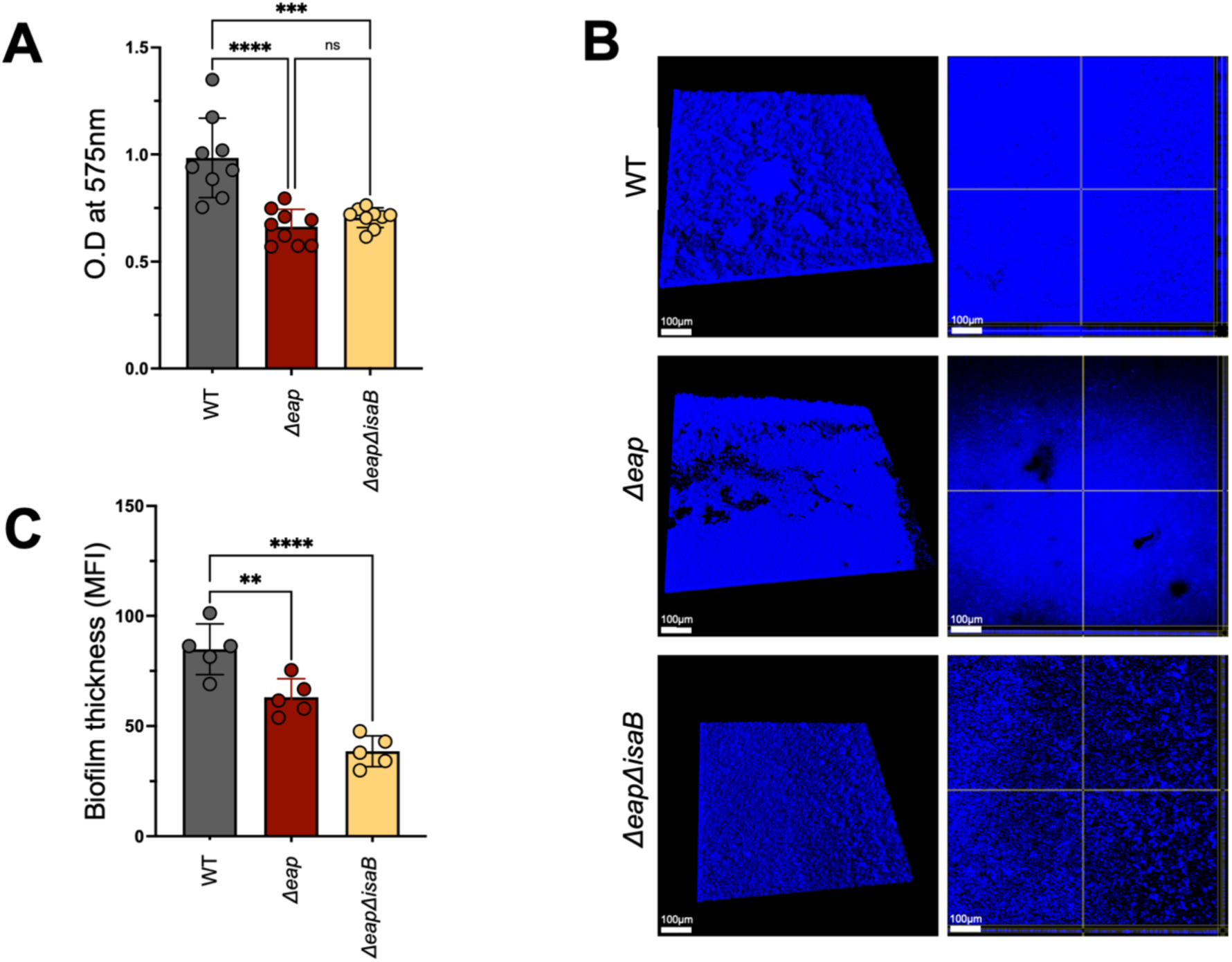
Eap proteins contribute to biofilm biomass. Crystal violet assay measuring biomass of WT, Δ*eap* and Δ*eapisaB* biofilms grown overnight in 96-well plates. Crystal violet staining was measured as O.D. at 575nm using previously established methods (A). 3-dimensional confocal microscopy images of WT, Δ*eap* and Δ*eapisaB* biofilms grown similarly to A, in 8-well chamber slides. Bacteria were stained with Hoechst Blue 33342 and images were captured at 400X magnification (left). Images of sections were taken close to the bottom of the biofilm for each strain (right) (B). Volume quantification of biofilms grown as described for B (C). Data represent 3 independent experiments performed in triplicate. Multiple comparisons were made with one-way analysis of variance and Tukey’s post hoc test. ****, P < 0.0001; ***, P < 0.001; **, P<0.01; ns, not significant. Images were taken using Imaris software. MFI calculations were done using ImageJ software.

### Eap proteins contribute to the porosity of *S. aureus* biofilms

To further investigate a role for Eap proteins in providing a specific structural advantage to *S. aureus* biofilms, we tested for differences in porosity when WT biofilms were compared to those formed by the isogenic mutants, *Δeap* and *ΔeapΔisaB.* We utilized 3 sizes of fluorescein isothio<u>c</u>yanate (FITC) labelled dextran (10k, 70k and 150k) and allowed biofilms to grow on 0.45µm membranes before measuring the levels of each FITC-dextran that could penetrate through biofilms formed by each strain using previously established methods [33]. While there were no differences between strains in the levels of 10k FITC-labelled dextran that could penetrate through biofilms (**Figure 2A**), we observed a significant increase in the porosity of mutants lacking Eap proteins compared to WT, when biofilms were incubated with 70 (**Figure 2B**) and 150k FITC-labelled dextran (**Figure 2C**). Additionally, we used 24-hour biofilms grown in 6-channel ibidi flow cells to image the entry and retention of various sizes of FITC-dextran as described above. Confocal images of biofilms incubated with each FITC-dextran for 1 hour, followed by 3 washes in saline revealed, that the levels of 70k and 150k (but not 10k) FITC-labelled dextran that could penetrate and be retained in *Δeap* and *ΔeapΔisaB* biofilms was higher than the WT control (**Figure 2D**). These data together indicate that Eap proteins reduce the overall porosity of *S. aureus* biofilms, and that the absence of these 3 proteins increases access to the biofilm.

**Figure 2.**
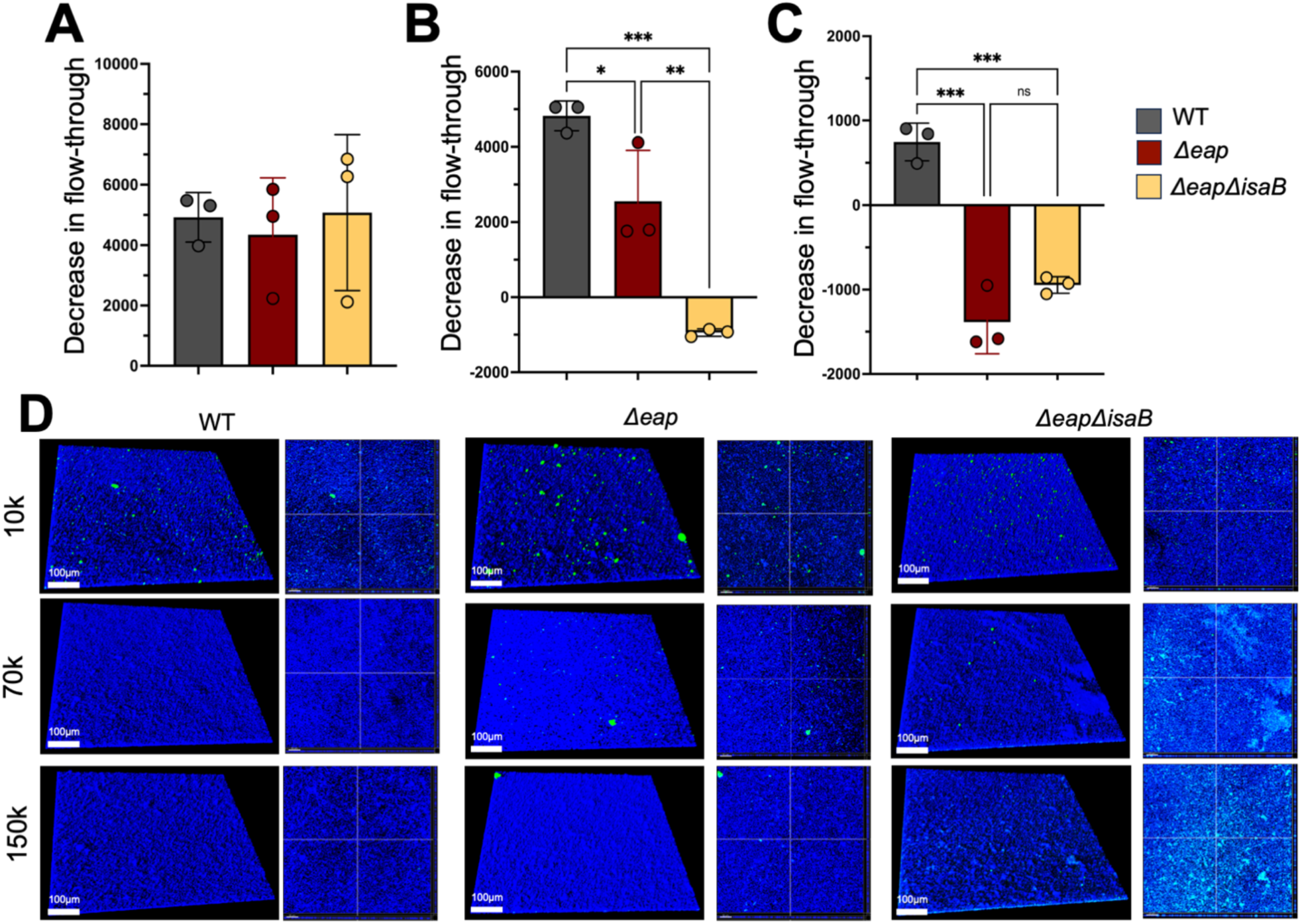
Eap proteins reduce biofilm porosity. Flow through of 10, 70 and 150kD FITC-labelled dextran from WT, Δ*eap* and Δ*eapisaB* biofilms grown on 0.45μm PVDF membranes. Results are calculated in relative fluorescence units as compared to control biofilms lacking dextran (A-C). Representative 3D confocal images of biofilms grown in 6-well ibidi chamber slides after incubation with FITC-labelled dextran of respective molecular weights (D). Multiple comparisons were made with one-way analysis of variance and Tukey’s post hoc test. ***, P < 0.001; **, P<0.01; *, P<0.1; ns, not significant. Images were taken using Imaris software.

### Eap proteins reduce macrophage invasion and phagocytosis of *S. aureus* biofilms

Previous reports describe a role for Eap in protecting planktonic *S. aureus* against human neutrophils. While Eap, EapH1 and EapH2 were shown to inhibit neutrophil proteases, Eap was shown to bind to neutrophil DNA and interfere with <u>n</u>eutrophil <u>e</u>xtracellular trap (NET) formation [24], [34]. Since we found that *S. aureus* biofilms lacking Eap proteins were significantly more porous with reduced biomass, we examined whether these differences would affect the ability of biofilms to evade phagocytosis by innate immune cells [27], [35]. Macrophages and neutrophils are crucial in the innate immune response to infection [36], [37], [38]. We therefore incubated mature biofilms with either primary murine bone marrow-derived macrophages or neutrophils for 4-6 h to quantify the number of leukocytes that could penetrate and phagocytose either WT or *Δeap* biofilms. Since no significant differences were observed between the *Δeap* and *ΔeapΔisaB* biofilms in earlier studies, WT was compared with the *Δeap* strain for these assays. Macrophage invasion into *Δeap* biofilms was significantly increased compared to WT, as reflected by both visualization (**Figure 3A, B**) and quantification (**Figure 3C**). Additionally, quantification of macrophages containing bacteria were also significantly higher in *Δeap* mutant biofilms compared to WT (**Figure 3D**) [27]. Lastly, the total number of observable macrophages associated with *Δeap* mutant biofilms was significantly higher compared to WT indicating that there were more intact macrophages that are phagocytosing bacteria in *Δeap* biofilms (**Figure 3E**). These results together demonstrate that Eap proteins reduce the invasion and phagocytosis of *S. aureus* biofilms by macrophages.

**Figure 3.**
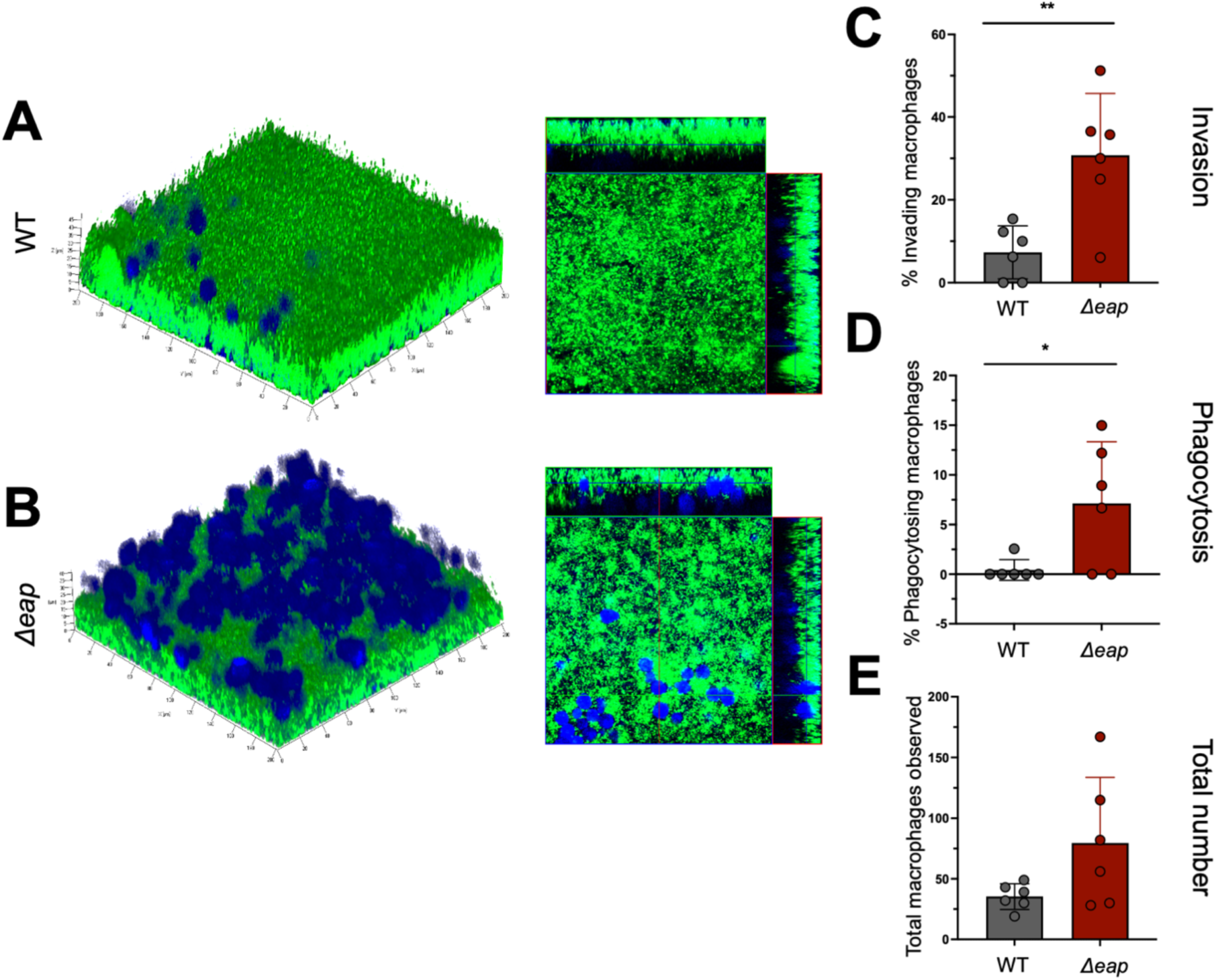
Eap proteins protect biofilms against macrophages. Representative 3D confocal images of green fluorescent protein (GFP)-labelled WT (A) or Δ*eap* (B) biofilms incubated with Cell Tracker Blue labelled macrophages for 4-6 hours (left). Cross sectional images from biofilms shown on left (right). Quantification macrophages invading WT or Δ*eap* biofilms (C), phagocytosing bacteria (D) and total numbers observed (E). Student *t* tests were performed for pairwise comparisons.**, P value = 0.0055; *, P value= 0.0261.

When similar experiments were performed with primary murine neutrophils, although larger numbers of neutrophils could be observed invading *Δeap* biofilms compared to WT (**Figure S1A and B**) this did not reach statistical significance (**Figure S1C**). Furthermore, there were no differences in the numbers of neutrophils observed phagocytosing biofilm (**Figure S1D**) or total number of neutrophils present (**Figure S1E**), between WT and *Δeap* biofilms. Collectively, these results indicate a larger role for Eap in preventing phagocytosis and clearance of biofilms by macrophages, in comparison to neutrophils.

### Eap proteins contribute to *S. aureus* prosthetic joint infection

Since biofilms lacking Eap proteins were more susceptible to invasion and phagocytosis by macrophages *in vitro* and exhibited less structural organization, we next examined whether these phenotypes would translate to altered biofilm survival *in vivo*. A previously established mouse model of prosthetic joint infection was used to compare the ability of *Δeap* to form biofilm compared to WT bacteria [39], [40]. Three time points were selected to reflect planktonic growth (day 3), transition to biofilm formation (day 7), and chronicity (day 14) based on recalcitrance to systemic antibiotics [29]. A larger number of animals was analyzed at day 7 since this represents the transition period to biofilm growth and was considered the best interval to interrogate potential phenotypes given the biofilm structural defects observed with *Δeap in vitro*. Bacterial burden was significantly reduced in the tissue surrounding the infected joint with *Δeap* bacteria at days 7 and 14 post-infection, which extended to the joint at day 7 with a trending decrease at day 14 (**Figure 4A-B**). Titers in the femur were also lower at days 7 and 14 with *Δeap*, although this did not reach statistical significance, and no differences were observed on the implant (**Figure 4C-D**). Previous work has described the role of <u>g</u>ranulocytic <u>m</u>yeloid-<u>d</u>erived <u>s</u>uppressor <u>c</u>ells (G-MDSCs) in promoting *S. aureus* biofilm survival by their ability to inhibit macrophage proinflammatory activity, neutrophil antimicrobial activity, and T cell activation [26], [39], [41]. Therefore, flow cytometry was performed on infected tissue samples to quantify G-MDSC infiltrates between WT and *Δeap* infected mice [42]. Although the overall number of CD45^+^ leukocytes trended higher in WT infected mice compared to those infected with *Δeap* bacteria (**Figure S2A**), G-MDSC infiltrates (CD45^+^Ly6G^+^Ly6C^+^) were similar between the groups (**Figure S2B**). Since neutrophils are also recruited to infected tissues, we measured the number of neutrophils (CD45^+^Ly6G^+^Ly6C^-^) in these animals (**Figure S2C**). While *Δeap* infected mice had lower neutrophil numbers compared to WT at day 7, these differences were not statistically significant at day 14. Altogether these data suggest that Eap proteins play specific roles in promoting *S. aureus* survival during biofilm-associated infection *in vivo.* Furthermore, while G-MDSC and neutrophil responses were generally comparable between WT and *Δeap* infected conditions, the consequence of this response to bacterial survival is likely altered as a function of Eap expression based on our *in vitro* findings.

**Figure 4.**
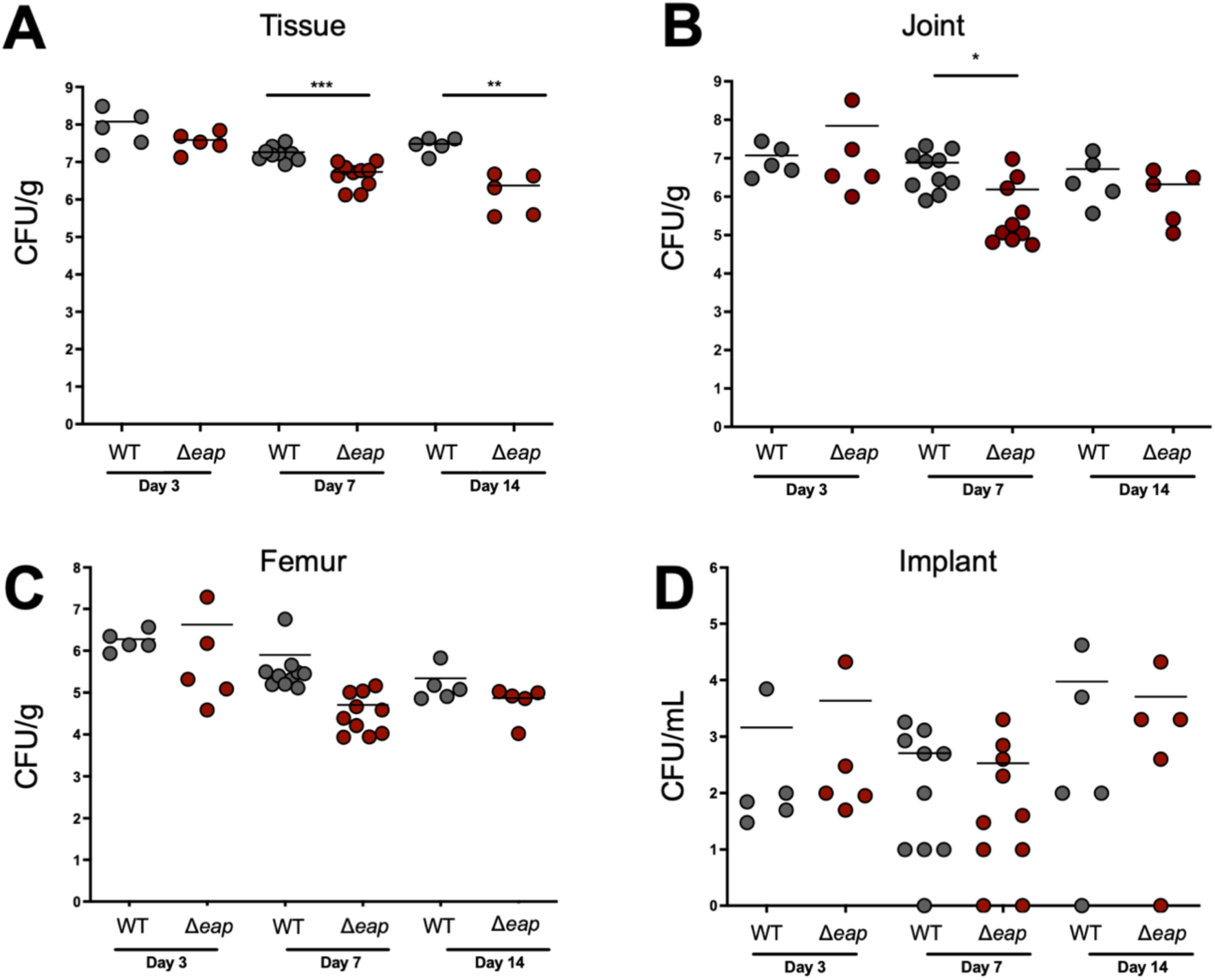
Eap proteins promote bacterial survival during prosthetic joint infection. Bacterial burdens were quantified in C57BL/6 mice infected with WT or Δ*eap S. aureus* at the indicated time points post-infection in the soft tissue surrounding the knee (A), joint (B), femur (C), and sonicated titanium implant (D)(n=5-10 mice/group). Student’s test was performed for pair-wise comparison.*** P value= 0.0001, ** P value= 0.0061 *P value= 0.0239.

## Discussion

This work provides evidence that Eap proteins are important to *S. aureus* biofilm structure and can influence the host response to infection. We demonstrate that Eap proteins increase the thickness (**Figure 1**) and reduce the porosity (**Figure 2**) of *S. aureus* biofilms, and this affects macrophage functions (**Figure 3**). Although to a smaller extent these proteins also confer an advantage to biofilms that are exposed to neutrophils (**Figure S1**). *In vivo,* while Eap proteins do not seem to alter the overall innate immune response to *S. aureus* biofilm infection, bacterial survival was significantly reduced with *Δeap* compared to WT bacteria (**Figure 4**). When taken together with our *in vitro* findings, this suggests that Eap proteins may serve to prevent bacterial clearance by phagocytes *in vivo* (**Figure 5**). It is unlikely that these phenotypes are due solely to the increased access afforded to phagocytic cells by virtue of a reduction in biofilm thickness and porosity.

**Figure 5.**
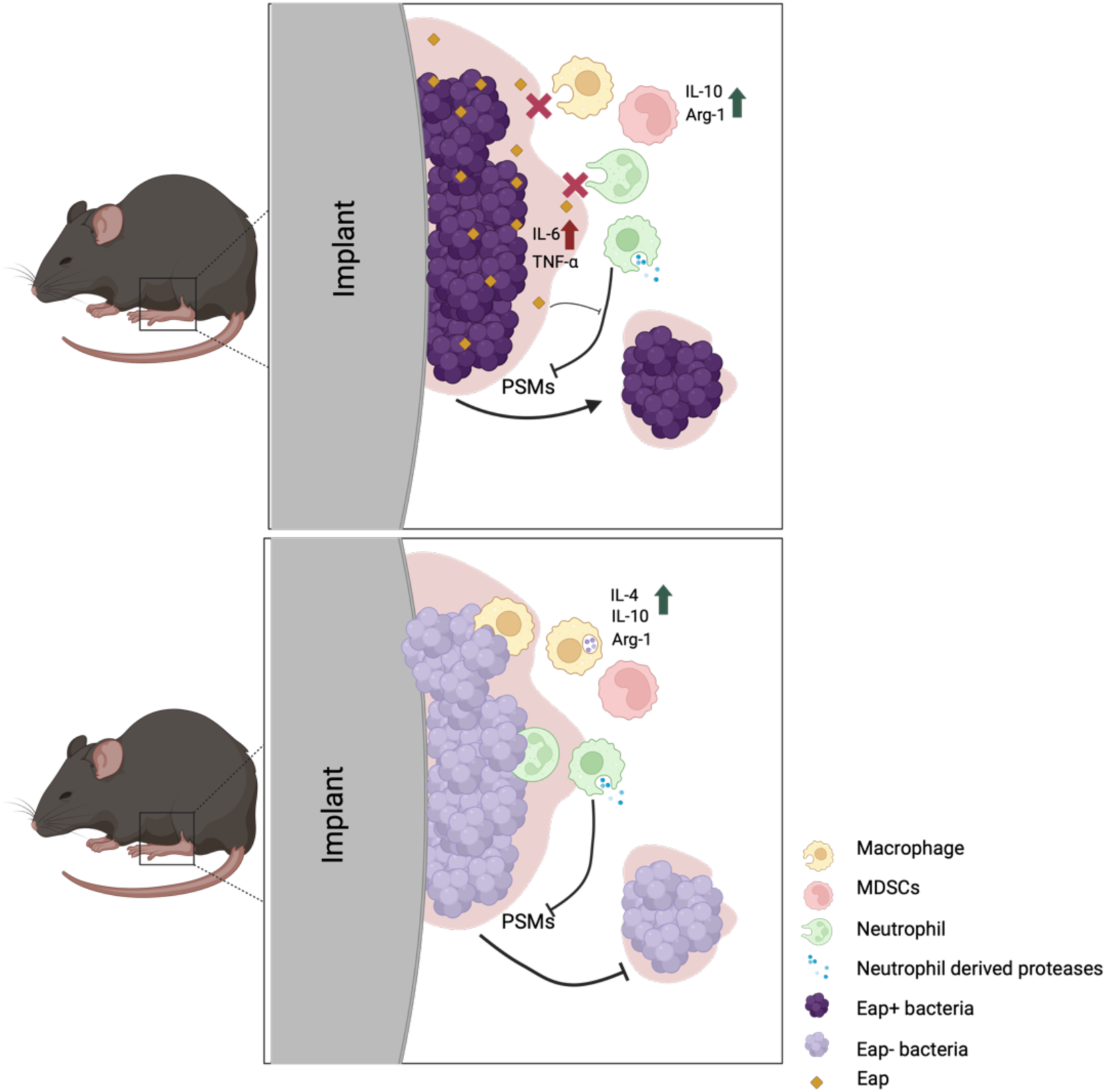
Summary and hypotheses based on current findings and previous literature. Expression of Eap protects biofilms from macrophage invasion and phagocytosis and, to a lesser extent, neutrophils. Proinflammatory signatures associated with Eap expression are likely dampened by the anti-inflammatory macrophage response to biofilms. Eap proteins prevent protease-mediated degradation of phenol soluble modulins (PSMs), which allows for subpopulations of the community to disperse and spread (A). In the absence of Eap, phagocytes gain some entry into the biofilm and likely phagocytose and kill bacteria. Proteases released from neutrophils degrade PSMs and prevent dispersion allowing the immune response to continue clearance of the biofilm infection. Inflammation is reduced in the presence of anti-inflammatory signatures generated from biofilm-exposed macrophages (B).

Previous reports have shown that Eap proteins are anti-inflammatory and immunomodulatory, with anti-protease activity specifically against human neutrophils. Eap is also known to prevent the degradation of phenol soluble modulin toxins (PSMs) by neutrophil-derived proteases [43]. PSMs can lyse neutrophils and are released during the transition of biofilms to planktonic growth, making them an important virulence factor during infection [44]. PSMs are also reported to form amyloid fibers that can stabilize biofilm structure [45]. Whether PSMs contribute to Eap-associated tolerance of neutrophils warrants further investigation. Similarly, while studies report a role for Eap in binding to DNA and blocking NET formation, *S. aureus* biofilms are documented to induce NETosis in a leukocidin-dependent manner and to utilize a nuclease to degrade the DNA released from neutrophils [34], [35], [46]. Since experiments with Eap were performed with purified protein and chemically induced NETs, analyzing the effect of Eap proteins on NETs released from biofilms would provide more information on the role of Eap proteins during neutrophil ET release [34]. Additionally, it has been demonstrated that once Eap binds DNA it does not cleave it [34]. It is therefore possible that Eap-bound host DNA can act as an immune evasion strategy *in vivo,* allowing *S. aureus* to appear as a ‘self’ molecule to the immune system, although this remains speculative.

Here we demonstrate that Eap proteins can provide *S. aureus* with some degree of protection against macrophages *in vitro*. Eap expressed by planktonic bacteria interacts with peripheral blood mononuclear cells presumably via the intercellular adhesion molecule 1 (ICAM-1) to induce proinflammatory cytokine production (IL-6, TNF-α) [47]. Biofilms bias macrophage towards an anti-inflammatory phenotype (arginase-1, IL-4, IL-10) that is compounded by the action of immune suppressive G-MDSCs known to impair T cell activation [27], [48]. These biofilm-specific mechanisms of macrophage subversion may therefore neutralize any proinflammatory signals generated as a function of Eap. Conversely, a number of reports provide evidence that Eap impairs neutrophil and T cell recruitment as well as T cell activation. These functions were attributed to higher concentrations of Eap such as those that would be produced by bacterial biofilms [21], [49], [50]. It is therefore likely that the anti-inflammatory properties of Eap are more relevant during biofilm infections, whereas proinflammatory processes are associated with survival of planktonic populations.

Lastly, in addition to DNA, Eap proteins have been documented to promiscuously bind multiple host-associated ligands including fibrinogen and collagen. Synovial fluid is an ultrafiltrate of blood plasma that encases joints and periprosthetic implants [51], [52], [53]. Whether Eap promiscuously binds to components of synovial fluid is currently unknown. This viscous fluid is known to harbor *S. aureus* aggregate biofilms reported to bind fibrinogen via its two sortase anchored fibronectin binding proteins FnbpA and B [54]. The properties of these biofilms are distinct from their surface-associated counterparts and can be formed by subpopulations of detached biofilm bacteria [55]. It is therefore plausible that the Eap proteins, FnbpA and FnbpB could contribute to biofilm survival at different phases of the infection lifecycle and require the additional activities of dispersion cues including PSMs to evade the immune response during prosthetic joint infection. Altogether this works builds on previous studies and adds to our knowledge of the innate immune response to *S. aureus* biofilm infections. **Figure 5** summarizes our findings and hypotheses based on current and previous work to depict how Eap proteins may be playing a multifactorial role during *S. aureus* biofilm-associated prosthetic joint infection, with potential new avenues of investigation to better understand the complex dynamics that make *S. aureus* a successful biofilm pathogen.

## Resource availability

### Lead Contact

Further inquiries and information on reagents and resources should be directed to (and will be fulfilled by) the lead contact, Alexander R. Horswill.

### Materials availability

Reagents and materials used or generated in this study can be made available upon request from the lead contact.

### Data availability

Data reported in this manuscript will be made available by the lead contact upon request.

## Author Contributions

Conceptualization, M.B., T.D.S., J.L., T. K., A.R.H.; Methodology, M.B., T.D.S., T. K., A.R.H; ; Investigation, M.B., T.D.S., J.L.; ; Writing-Original Draft, M.B., Writing-Review and Editing, M.B., T.K., A.R.H.; Funding Acquisition, T.K, and A.R.H.; Supervision, T.K., A.R.H.

## Acknowledgments

The authors would like to thank members of the Horswill and Kielian groups for their critical evaluation of the data in this manuscript. This work was funded by the NIH/NIAID grant(s) AI083211 to A.R.H. and T.K., and VA Merit Award BX002711 to A.R.H

## Declaration of interests

The authors declare no competing interests.

## Supplemental Information

Document S1. Figures S1, S2

## Materials and Methods

### Bacterial strains and growth conditions

Unless otherwise indicated all experiments were performed in the USA300 clinical strain background. Bacterial cultures were grown in tryptic <u>s</u>oy <u>b</u>roth (TSB) at 37°C with shaking (200RPM).

### Construction of *S. aureus* bacterial mutants

Chromosomal deletions of the three Eap encoding genes (*eap, eapH1* and *eapH2*) were performed using previously established methods [33]. Briefly, the temperature sensitive pJB38 plasmid was used to introduce DNA fragments (∼1kb) flanking the target region of interest. Flanking DNA was amplified (Phusion high fidelity polymerase, NE Biolabs) using gene specific primers, products were digested with restriction enzymes (Table 2) and purified (Qiagen PCR purification). Following triple ligation into pJB38, the plasmid was electroporated into *E. coli* DC10b and selected for on Luria Bertani agar plates containing 100μg/mL ampicillin. Following confirmation from single colonies, plasmid was purified, PCR used for confirmation with construction and sequencing primers performed and plasmid was electroporated into S. aureus. Positive clones were selected on tryptic <u>s</u>oy <u>a</u>gar (TSA) containing 10μg/mL chloramphenicol and homologous recombination performed at 42 degrees for 24 hours. Following overnight incubation in TSA-Cam and a series of subcultures in TSB at 30 degrees, counterselection was performed on 200 ng/ml anhydrotetracycline (30 degrees/ overnight). Loss of plasmid was indicated by growth on TSA but not TSA-Cam and presence of desired mutations were verified using PCR with chromosomal primers that were outside the region of mutation.

### *In vitro* 24-hour biofilm growth

All *in vitro* biofilms used for biomass and matrix porosity measurements were grown in TSB containing 0.4% glucose as previously published, unless otherwise indicated [33], [56]. Bacterial cultures were grown overnight (16-18 hours) in TSB at 37°C with shaking (200RPM). The next day, bacteria were sub cultured (1:100) in fresh TSB for 2-3 hours and brough to exponential phase corresponding to an <u>o</u>ptical <u>d</u>ensity (OD) at 600nm of 0.5-0.7 as previously described. Cultures were then centrifuged at 3900RPM for 2 minutes, washed once with <u>p</u>hosphate <u>b</u>uffered <u>s</u>aline (PBS), centrifuged and re-suspended in TSB containing 0.4% glucose for biofilm growth measurements.

### Biofilm biomass measurements using crystal violet staining

Cultures were prepared in TSB containing 0.4% glucose as described above. Bacteria were seeded into 96-well microtiter plates (Costar, 200μL per well) and incubated overnight at 37°C in a humidified chamber for 24 hours. Biofilms were washed with double distilled water (dd water) and incubated with 0.1% crystal violet for 30 minutes at room temperature. Crystal violet was drained, and plate was washed in dd water 3 times followed by addition of 33% acetic acid to the wells. After a 30-minute incubation, solubilized biofilms were pipetted into a new 96 well plate and O.D was measured at 575nm. Measurements were made in comparison to well containing PBS.

### *In vitro* biofilms for confocal imaging

Cultures were prepared in TSB containing 0.4% glucose as described above. Bacteria were seeded into 8-well ibidi μ-slides (ibidi, Cat.No:80826) and incubated for 24 hours at 37°C in a humidified chamber. Spent media was removed, biofilms were washed with PBS and stained with 10μg/mL Hoechst Blue 33342 stain (Thermo Fisher, Cat.No: H3570) for 30 minutes for confocal imaging. Biofilms were then washed again with PBS and fixed with 10% formalin. Biofilms were visualized using the Olympus FV1000 confocal laser scanning microscope using the Z-stack feature to collect 3D images spanning the thickness of the biofilm. All experiments were performed with 2 technical duplicate biofilms per strain for a total n= 4 (n=8 biofilm technical replicates per strain). 3 images were taken per technical biofilm replicate (n=24 images per strain).

### Measuring porosity of *in vitro* biofilms

24-hour biofilms of WT or respective isogenic mutants were grown as described above in 96 well plates containing 0.45μm PVDF membranes as previously described [33]. Briefly, biofilms grown in 96-well plates without a membrane were used as a negative control. Following 24-hour growth, control biofilm biomass was measured using the crystal violet assay described above. Media was removed from filter plates and replaced with 100μL MES (2-(N-<u>m</u>orpholino)<u>e</u>thane <u>s</u>ulfonic acid) (MES) buffer containing 1mg/mL FITC-isocyanide-dextran (with dextran at a molecular weight of either 4K, 10K, 70K, or 150K). These experiments were performed using a negative control consisting of biofilms resuspended in buffer lacking FITC-dextran. Filter plates were centrifuged for 45 seconds at 20g, flow through collected and relative levels of fluorescence measured with excitation and emission wavelengths of 470 nm and 523 nm respectively. Values were plotted in comparison to the media-only control as a measure of maximum fluorescence. For microscopy, bacteria were grown as described above and seeded into 8-well ibidi μ-slides (ibidi, Cat.No:80826). Biofilms were washed in PBS and resuspended in 1mg/mL FITC-isocyanide-dextran of various sizes as described above, for 1 hour. 3D images were taken using the Olympus FV1000 system.

### *S. aureus* biofilm-leukocyte co-culture experiments

Confocal microscopy experiments depicting the interaction of macrophages or neutrophils with *S. aureus* biofilms were performed as previously published [27]. Briefly, green fluorescent protein (GFP) labelled bacteria were grown to exponential phase as described above and seeded into chamber slides coated with human plasma. Biofilms were allowed to grow for 4 days at 37°C and incubated with Cell Tracker Blue labelled bone marrow-derived macrophages or thioglycolate-elicited neutrophils from C57BL/6 mice for 4-6 h using a Zeiss laser scanning confocal microscope (LSM 710 META; Carl Zeiss). 3D images of biofilms were collected using Xen 2007 software (Carl Zeiss) as previously described [27]. The number of leukocytes invading and phagocytosing biofilms was quantified by measuring the distance of immune cells from the biofilm base (invasion) and number of leukocytes with intracellular bacteria in each field of view using orthogonal images.

### Mouse prosthetic joint infection model

*S. aureus* biofilm infection was studied *in vivo* using an established model of implant-associated prosthetic joint infection. Briefly, 8–10-week-old male and female C57BL/6 mice (n=5-10 mice/time point/strain) were used to introduce an implant into the intramedullary canal of the femur as previously described [26], [30], [39], [42]. Approximately 10^3^ of WT or *Δeap* bacteria were inoculated at the implant tip and animals were administered buprenorphine slow release (SR) after surgery for pain relief. Animals were euthanized at days 3, 7, and 14 post-infection to collect tissue and implant samples as previously described [42]. Tissue homogenates and sonicated implants were plated on TSA containing 5% sheep blood to quantify total colony forming units (cfu) per gram of tissue or per mL diluent for implants. The soft tissue surrounding the knee was collected to quantify G-MDSC and PMN infiltrates by flow cytometry using antibodies for CD45, Ly6G, and Ly6C as previously described [42].

## Notes

### Competing Interest Statement

The authors have declared no competing interest.

