## Supplemental Figures for "Matrix porosity is associated with *Staphylococcus aureus* biofilm survival during prosthetic joint infection"

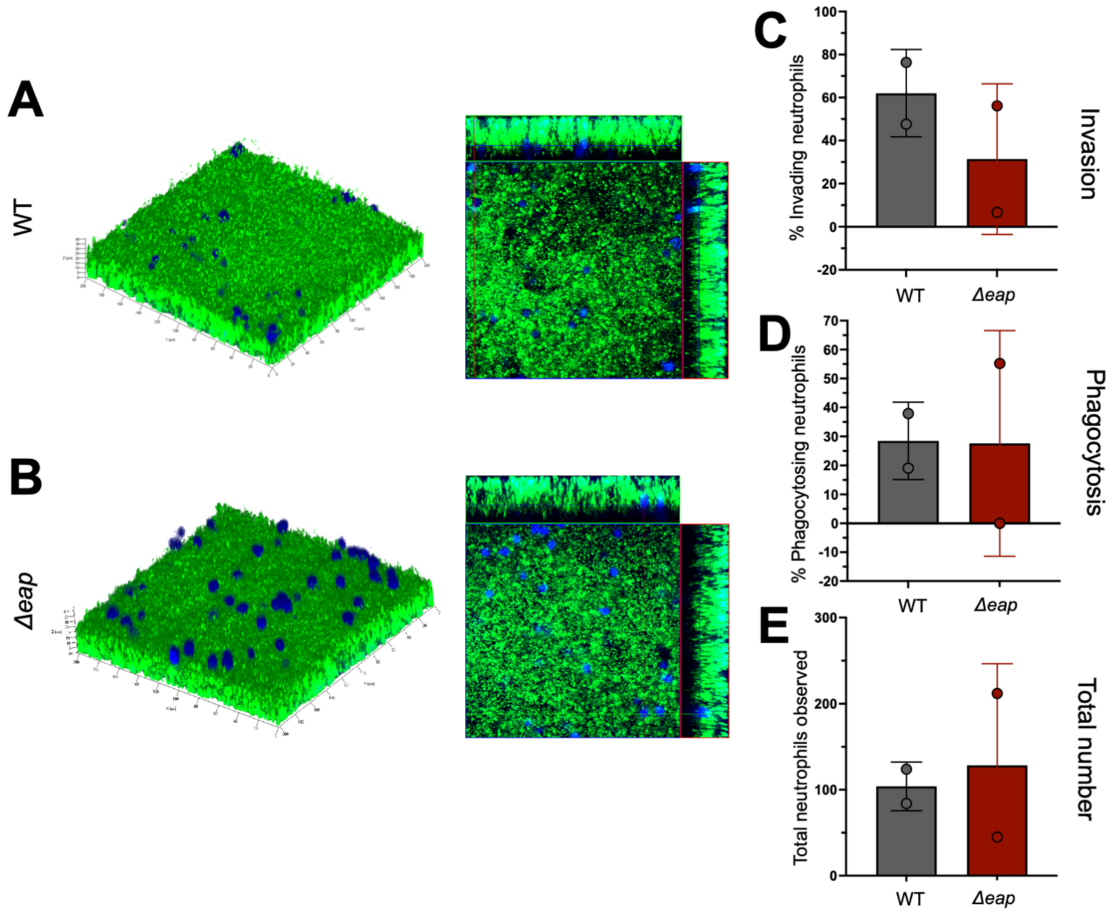


**Figure S1. Eap proteins do not influence neutrophil responses to biofilm.** Representative 3D confocal images of green fluorescent protein (GFP)-labelled WT (A) or Δ*eap* (B) biofilms incubated with Cell Tracker Blue labelled neutrophils for 4-6 hours (left). Cross sectional images from biofilms shown on left (right). Quantification of neutrophils invading WT or Δ*eap* biofilms (C), phagocytosing bacteria (D) and total numbers observed (E). Student's test was performed for pair-wise comparison.

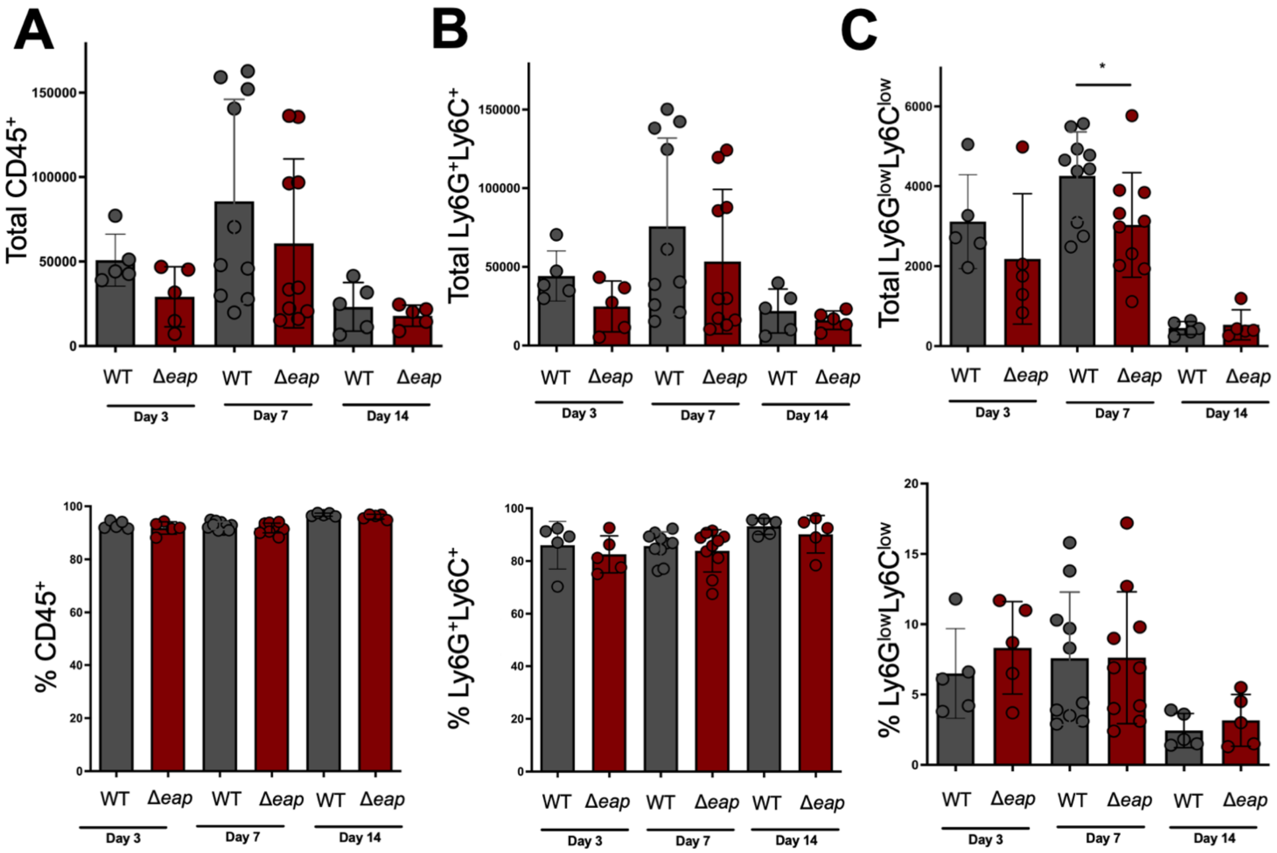


**Figure S2. Eap proteins do not affect granulocyte recruitment *in vivo.*** Flow cytometry quantification of $CD45^+$ populations, granulocytic myeloid-derived suppressor cell (defined as $CD45^+Ly6G^+Ly6C^+$) (B) and neutrophil (defined as CD45+ $Ly6C^{low}$ $Ly6G^{low}$) (C) populations in tissue homogenates of animals infected with either WT or Δ*eap* bacteria at days 3, 7 and 14 post-infection. Data is presented as total numbers and percentage of each total population. Student's test was performed for pair-wise comparison. *P value= 0.0366
